# Characterization of a clinical *Enterobacter hormaechei* strain belonging to epidemic clone ST418 co-carrying *bla*_NDM-1_, *bla*_IMP-4_ and mcr-9.1

**DOI:** 10.1101/2020.09.25.314500

**Authors:** Wei Chen, Zhiliang Hu, Shiwei Wang, Doudou Huang, Weixiao Wang, Xiaoli Cao, Kai Zhou

**Author notes:** To whom correspondence should be addressed: Dr. Xiaoli Cao, and Dr. Kai Zhou. Contributed equally to this study (Wei Chen, Zhiliang Hu).

## Abstract

An *Enterobacter hormaechei* isolate (ECL-90) simultaneously harboring *bla*_NDM-1_, *bla*_IMP-4_ and *mcr-9*.*1* was recovered from the secretion specimen of a 24-year-old male patient in a tertiary hospital in China. The whole genome sequencing of this isolate was complete, and 4 circular plasmids with variable sizes were detected. MLST analysis assigned the isolate to ST418, known as a carbapenemase-producing epidemic clone in China. *bla*_IMP-4_ and *mcr-9*.*1* genes were co-carried on an IncHI2/2A plasmid (pECL-90-2) and *bla*_NDM-1_ was harbored by an IncX3 plasmid (pECL-90-3). The genetic context of *mcr-9*.*1* was identified as a prevalent structure, “*rcnR-rcnA-pcoE-pcoS*-IS*903*-*mcr-9-wbuC*”, which is a relatively unitary model involved in the mobilization of *mcr-9*. Meanwhile, *bla*_NDM-1_ gene was detected within a globally widespread structure known as NDM-GE-U.S (“IS*Aba125*– *bla*_NDM-1_–*bla*_MBL_”). Our study warrants that the convergence of genes mediating resistance to last-resort antibiotics in epidemic clones would largely facilitate their widespread in clinical settings, thus representing a potential challenge to clinical treatment and public health.

**Importance:** Carbapenemase-producing *Enterobacteriaceae* (CPE) spread at a high rate and colistin is the last-resort therapeutic for the infection caused by CPE. However, the emergence of plasmid-borne *mcr* genes highly facilitates the wide dissemination of colistin resistance, thus largely threatens the clinical use of colistin. Here, we for the first time characterized a clinical *Enterobacter hormaechei* strain co-producing *bla*_NDM-1_, *bla*_IMP-4_ and *mcr*-9.1 belonging to an epidemic clone (ST418). The accumulation of genes mediating resistance to last-resort antibiotics in epidemic clones would largely facilitate their widespread in clinical settings, which may cause disastrous consequence with respect to antimicrobial resistance. Understanding how resistance genes were accumulated in a single strain could help us to track the evolutionary trajectory of drug resistance. Our finding highlights the importance of surveillance on the epidemic potential of colistin-resistant CPE, and effective infection control measures to prevent the resistance dissemination.

---

The worldwide prevalence of carbapenem-resistant *Enterobacteraceae* (CRE) has been well known (1), with *Klebsiella pneumoniae* carbapenemases (KPCs), New Delhi metallo-lactamases (NDMs), imipenemases (IMPs) and OXA-48-like enzymes being the most prevalent carbapenemase (2). Moreover, co-occurrence of these genes in a single strain has been frequently identified (3, 4). Currently, colistin is one of the last-resort antibiotics for the CRE infections (5). However, the emergence of plasmid-borne colistin resistance gene *mcr* frustrates colistin’s efficiency. MCR encodes a phosphoethanolamine transferase which is involved in the modification of lipopolysaccharide, the target of colistin. At present, the prevalence of *mcr* is the most widespread mechanism of colistin resistance (6). As of now, ten different *mcr* variants noted from *mcr*-*1* to *mcr-10* have been described (6-8). Of more concern, co-existence of *mcr* and carbapenemase genes has been sporadically reported (9). For instance, *mcr-1, mcr-3*.*5*, and *bla*_NDM-5_ are found in an *Escherichia coli* isolate (10), *mcr-4*.*3* and *bla*_NDM-1_ are co-identified in a clinical *E. cloacae* isolate from China (11), and *bla*_VIM-4_ and *mcr*-9 are found in an *E. hormaechei* isolate in USA (12). The rapid emergence of such new resistance phenotypes has broken through the last defense line and severely limited therapeutic options. In this report, we characterized a clinical *E. hormaechei* isolate simultaneously harboring *bla*_NDM-1_, *bla*_IMP-4_ and *mcr*-9.1. The structures of plasmids carrying three resistance genes were fully dissected to understand their dissemination and accumulation pattern.

This *E. hormaechei* isolate (ECL-90) was recovered from the secretion specimen of a 24-year-old male patient in October, 2017, who was admitted into a large tertiary hospital in Nanjing because of an acute community-acquired pneumonia. The patient subsequently developed a left orbital cellulitis resulting from the ascending infection of the respiratory tract. After receiving endoscopic sinus opening and drainage, and an anti-infective regimen including meropenem, tigecycline and fosfomycin, the patient recovered and was therefore discharged.

ECL-90 was first identified as *E. cloacae* complex by matrix-associated laser desorption ionization-time of flight mass spectrometry (BioMerieux, Craponne, France), and further confirmed as *E. hormaechei* by whole genome sequencing (WGS). Antimicrobial susceptibility test using microbroth dilution method showed that this strain was resistant to ertapenem (16 μg/ml), imipenem (8 μg/ml), meropenem (>16 μg/ml), cefepime (>32 μg/ml), ceftazidime (>32 μg/ml), cefotaxime (>32 μg/ml), cefuroxime (>64 μg/ml), cefazolin (>32 μg/ml), cefmetazole (>64 μg/ml), piperacillin/tazobactam (>256 μg/ml), amikacin (8 μg/ml), gentamicin (64 μg/ml), funantuoyin (128 μg/ml), trimethoprim and sulphame-thoxazole (>32 μg/ml), aztreonam (>128 μg/ml), piperacillin (>256 μg/ml), ciprofloxacin (8 μg/ml), levofloxacin (4 μg/ml), and ceftazidime/avibactam (>32 μg/ml), while remained susceptible to aztreonam/avibactam (<0.25 μg/ml), tigecycline (1 μg/ml), colistin B (0.25 μg/ml) and fosfomycin (32 μg/ml).

To better understand the resistance mechanism and genomic characterization of this strain, the genomic DNA was extracted and WGS was performed on the Hiseq 4000 instrument (Illumina, San Diego, CA, USA). Multi-locus sequence typing (MLST) analysis using MLST 2.0 assigned ECL-90 to sequence type (ST) 418 (allelic profile 53-35-154-44-45-4-6) (13), which is known as one of the predominant epidemic clones of carbapenemase-producing *E. cloacae* in China (14, 15). Antibiotic resistance genes were identified by using Resfinder v2.1 (http://cge.cbs.dtu.dk/services/ResFinder-2.1/), including genes conferring beta-lactam resistance (*bla*_NDM-1_, *bla*_IMP-4_, *bla*_SFO-1_, *bla*_SHV-12_, *bla*_TEM-1B_, *bla*_ACT-16_), colistin resistance (*mcr-9*.*1*), aminoglycoside resistance [*aac(3)-IId, aac(6’)-IIc, aac(6’)-Ib3, aph(3’’)-Ib, aph(3’)-Ia, aph(6)-Id, aph(6)-Id*], fluoroquinolone and aminoglycoside resistance [*aac(6’)-Ib-cr*], macrolide resistance (*ereA, mphA*), phenicol resistance *(catA2*), sulphonamide resistance (*sul1*), tetracycline resistance (*tetD*), and trimethoprim (*dfrA19*). Overall, the genotypes identified were consistent with the resistance phenotypes.

ECL-90 was susceptible to colistin although *mcr-9*.*1* was detected. The IPTG-induced expression of *mcr*-9.1 in *E. coli* BL21(DE3) containing pET28a-*mcr*-9.1 did not confer resistance to colistin, and the resistance to colistin could not be induced by using sub-MIC concentration of colistin. This is consistent with the previous report that *mcr* -9.1 is inactive in colistin resistance (16). A recent study with global data revealed that *Enterobacter spp*., was the predominant host of *mcr-9* (37%) (17). However, the underlying mechanism remains unclear. Broth conjugation assays using a sodium azide-resistant *E. coli* J53 isolate as a recipient showed that *bla*_NDM-1_ was transferable, but not for *mcr-9*.*1* and *bla*_IMP*-*4_.

S1-pulsed-field gel electrophoresis (S1-PFGE) analysis of the strain ECL-90 displayed 4 plasmids. To obtain full sequences of the plasmids, this isolate was further sequenced by Nanopore platform (Nanopore, Oxford, UK), and hybrid assembly was performed with Illumina sequencing data by using Unicycler version 0.4.8 (18), resulting in a chromosome with size of 4,584,517 bp, and 4 plasmids (pECL-90-1, pECL-90-2, pECL-90-3, and pECL-90-4) ranging in sizes from 6,364 bp to 348,891 bp (Table 1). The completeness of these plasmids was further verified by PCR loop experiment. *fosA* and *bla*ACT-16 were identified in the chromosome, and the other resistance determinants were carried by pECL-90-2 and pECL-90-3.

*bla*_IMP-4_ and *mcr*-9.1 were co-identified on an IncHI2/2A-type plasmid pECL-90-2, which was 348,891 bp in length with an average GC content of 48.59% (Fig.1). Consistently, a recent report identified that IncHI2-type plasmids may serve as a critical reservoir of *mcr-9* (17). Blasting the sequence of pECL-90-2 in GenBank database showed the best matches were plasmid pGW1 carried by a *Cronobacter sakazakii* strain GZcsf-1 (CP028975, 86.8% query coverage and 99.99% sequence identity) and plasmid p17277A_477 carried by a *Klebsiella quasipneumoniae subsp. quasipneumoniae* strain M17277 (CP043927, 79.54% query coverage and 99.99% sequence identity). This suggests that the plasmid could widely disseminate among *Enterobacteriaceae*. The genetic context of *mcr-9*.1 gene identified here was “*rcnR-rcnA-pcoE-pcoS*-IS*903*-*mcr-9-wbuC*” (Fig.1), which is known as a prevalent structure for *mcr-9* (17). Previous studies showed that a “IS*903*-*mcr-9-wbuC*-IS*26”* genetic structure has been found in 71% sequences harboring *mcr-9* in the NCBI Nucleotide Collection database (19), indicating the importance of IS*903B* in the spread of *mcr-9* gene. The downstream regulatory genes (*qseC* and *qseB*) detected in p17277A_477 (CP043927) were replaced by an IS*26* here (Figure 1).

**FIG 1.**
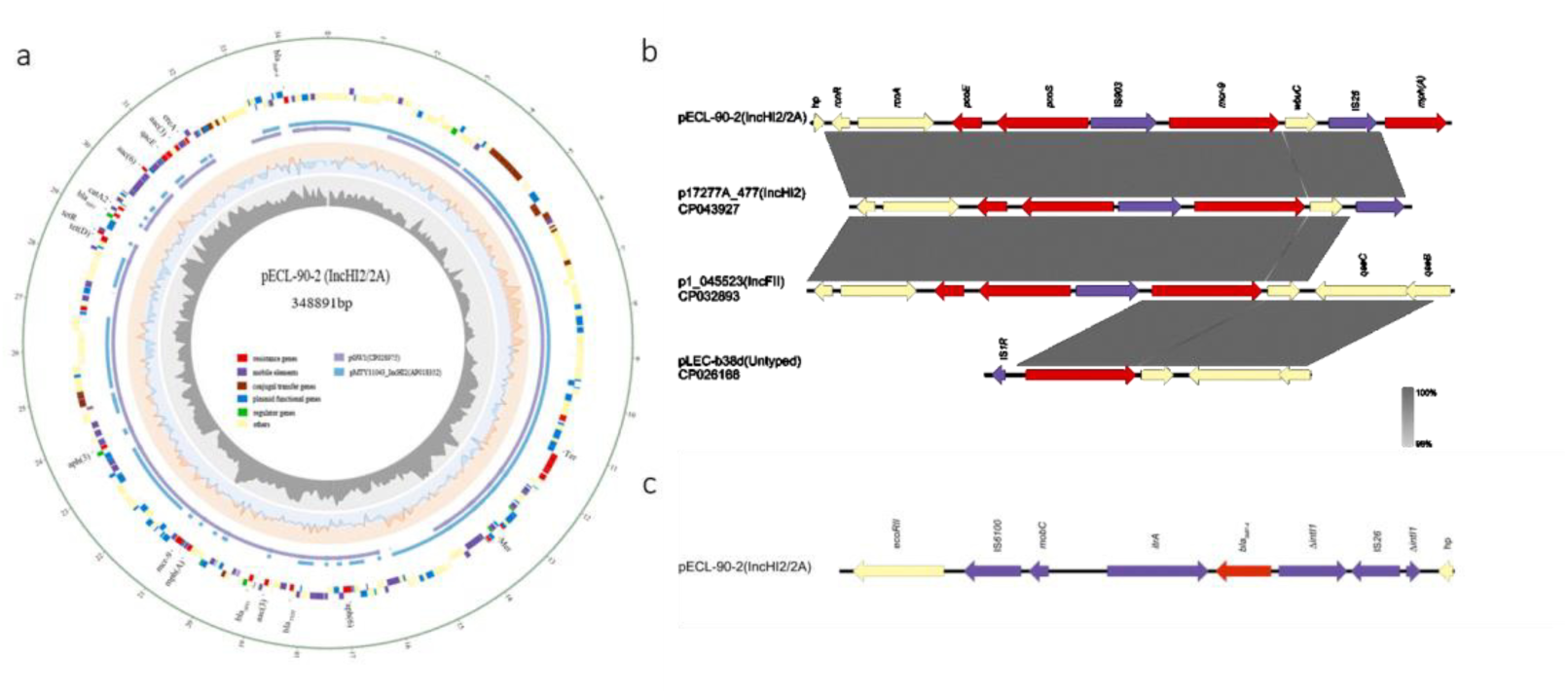
Analysis of *mcr-9.1*-harboring IncHI2/2A-type plasmid pECL-90-2 and the genetic context of *mcr-9.1* and *bla*_IMP-4_. (a) Plasmid structure of pECL-90-2 compared to pGW1 (GenBank accession number CP028975) and pMTY11043_IncHI2 (GenBank accession number AP018352) is shown. Open reading frames are indicated by colored columns based on predicted gene function. Dark gray, GC content; light blue, GC skew (+); orange, GC skew (-); (b) Comparison of mcr-9.1 genetic context harbored by pECL-90-2, p17277A_477 (IncHI2) (GenBank accession number CP043927), P1_045523 (IncFII) (GenBank accession number CP032893), and pLEC-b38d (Untypeable) (GenBank accession number CP026168). Dark fray shading denotes regions of shared homology among different plasmids; (c) Genetic context of *bla*_IMP-4_ carried by pECL-90-2.

The *bla*_IMP-4_ gene was carried by a class I integron designated as In*823b*, which located in an IS*6100*-IS*26* transposon-like structure (20, 21). The structure has previously been identified as being prevalent mediating the dissemination of *bla*_IMP-4_ gene in China (21). The *intl1* gene of In*823b* was disrupted by the insertion of IS*26*, and a single resistance gene cassette *bla*_IMP-4_-attC*bla*_IMP-4_ adjacent to a group IIc intron Kl.pn.I3 was identified (Fig.1). The typical 3’-conserved segment of In*823b* was absent. Altogether, numerous mobile genetic elements flanking *mcr-9.1* and *bla*_IMP-4_ gene indicate the transferable potential independent of the plasmid mobilization.

The *bla*_NDM-1_ gene was carried on a 44,961-bp IncX3 plasmid (pECL-90-3) (Fig.2). IncX3-type plasmids mediating the dissemination of *bla*_NDM-1_ among these homologous strains have been previously evidenced intensively (22). Query against GenBank showed that pECL-90-3 shared the highest similarity with pNDM5-L725 carried by *E. coli* strain L725 (CP036205, 99.73% query coverage and 99.99% sequence identity) and pBM527-2 carried by *Citrobacter sp*. strain CF971 (CP041048, 99.73% query coverage and 99.99% sequence identity). The genetic context of *bla*_NDM-1_ gene detected here is “IS*Aba125*–*bla*_NDM-1_–*ble*_MBL_” (Fig.2), which has recently been named NDM-GE-U.S. and has been found to be widespread globally (23). The wider context of *bla*_NDM-1_ gene IS*Aba125*–*bla*_NDM-1_–*ble*_MBL_-*ΔtrpF*-*dsbC* was flanked by IS*3000* and IS*26*, which was identical to that detected in *E. coli* strain BJ01 (JX296013) isolated in China (24).

**FIG 2.**
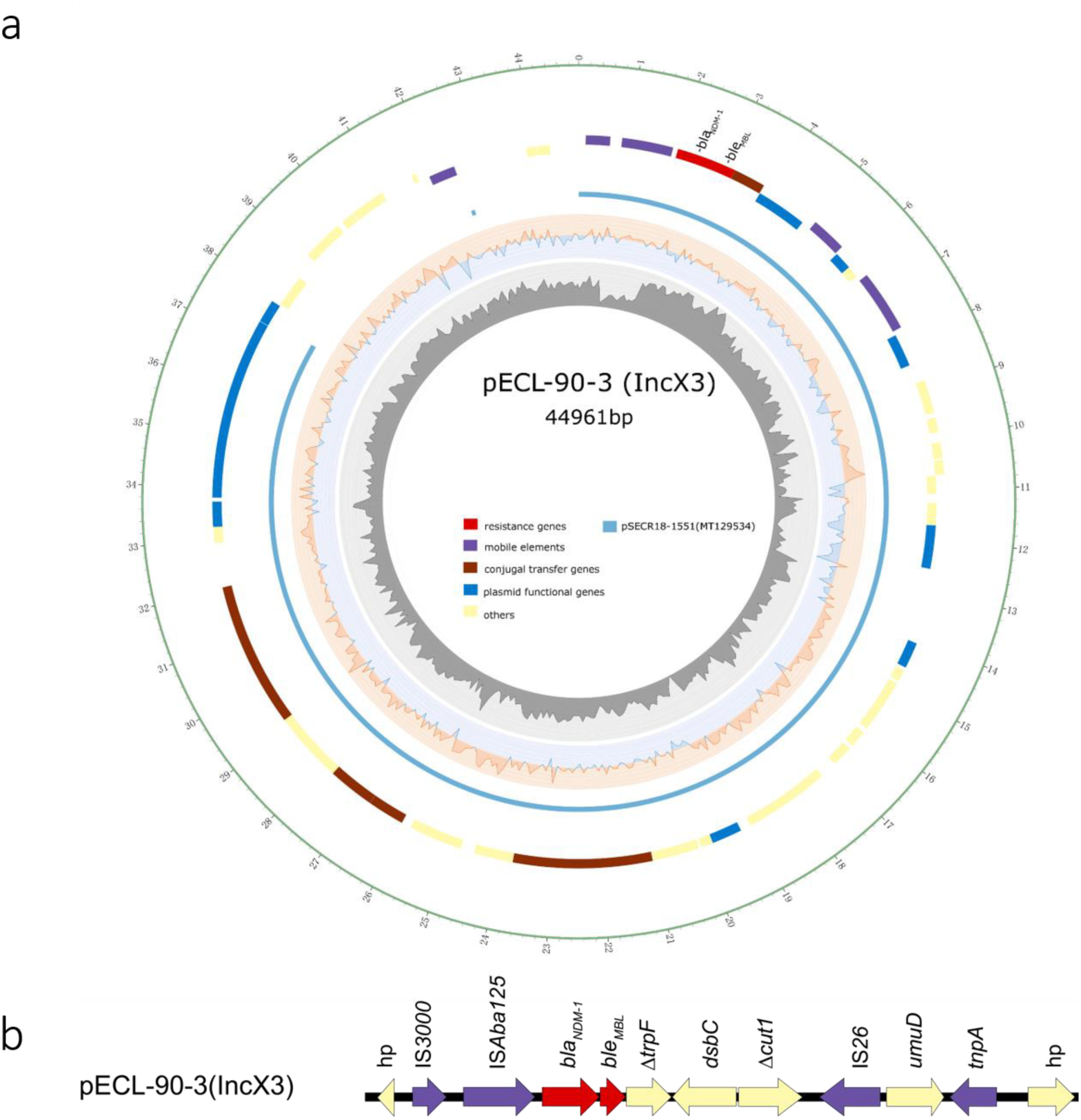
Analysis of *bla*_NDM-1_-harboring IncX3-type plasmid pECL-90-3 and the genetic context of *bla*_NDM-1_. (a) Plasmid structure of pECL-90-3 compared to pSECR18-1551 (GenBank accession number MT129534). Open reading frames are indicated by colored column based on predicted gene function. Dark gray, GC content; light blue, GC skew (+); orange, GC skew (-); (b) Genetic context of *bla*_NDM-1_ detected on pECL-90-3.

Noteworthily, the backbone structures of plasmids identified in our study shared a high similarity with plasmids harbored in multiple *Enterobacteriaceae* family such as *E. coli*, and *K. pneumoniae*. Under the selective pressure of antimicrobial agents, these resistance-encoding determinants might be recruited into a variable genetic locus flanked by mobile elements such transposons and insertion sequences, leading to a successful transmission among various *Enterobacteriaceae* species. Especially, the convergence of genes mediating the resistance to last-resort antibiotics (e.g. carbapenems and colistin) in epidemic clones would largely facilitate their widespread in clinical settings, thus represents a potential challenge to clinical treatment and public health.

In summary, we here for the first time characterized a clinical epidemic *E. hormaechei* clone ST418 co-harboring *bla*_NDM-1_, *bla*_IMP-4_ and *mcr*-9.1. The accumulation of genes conferring resistance to last-resort antibiotics via various mobile genetic elements highlights that stricter infection control measurements should be conducted to prevent the dissemination of such “chimera superbug”.

## Accession number (s)

The chromosome of ECL-990 has been deposited in GenBank nucleotide sequence database under accession number of CP061744; four plasmids have been deposited in the GenBank nucleotide sequence database under accession number of CP061745 (pECL-90-1), CP061746 (pECL-90-2), CP061747 (pECL-90-3), and CP061748 (pECL-90-4).

## Acknowledgments

This study was supported by the Nanjing Medical Science and technique Development Foundation (QRX17059), and National Natural Science Foundation of China (81902124 and 81702045).

